# Synergistic Cytotoxicity of Permethrin and N,N-Diethyl-Meta-Toluamide on Sinonasal Epithelial Cells

**DOI:** 10.1101/2024.08.27.610017

**Authors:** Jivianne T. Lee, Saroj K. Basak, Hong-Ho Yang, Kimberly A. Sullivan, Tom Maxim, Daniel S. Shin, Nancy Klimas, Eri S. Srivatsan

## Abstract

**Background:** N,N-Diethyl-Meta-Toluamide (DEET) and permethrin are pesticides commonly used in combination due to their synergistic insecticidal and repellent properties. This study investigates whether simultaneous exposure to these compounds elicits synergistic cytotoxicity in sinonasal epithelial cells (SNECs).

**Material and Methods:** Ethmoid sinus mucosal specimens were procured from eight patients during endoscopic sinus surgery. SNECs were expanded on culture plates and exposed to various concentrations of DEET and permethrin (0-5μm), individually and concurrently, for up to 156 hours. Experiments were replicated in triplicates and cell viability was recorded every 2 hours using *IncuCyte* real-time cell imaging system. Synergy score was calculated on the basis of Loewe additivity synergy finder model.

**Results:** DEET and permethrin exhibited synergistic cytotoxicity across all eight tissues, albeit with variations in onset and magnitude. Peak synergy was observed at 144h for tissue 1 (*S_Loewe_* 4.2, 95% CI 1.8-7.3; [permethrin concentration, DEET concentration] [2.5μ*M*, 1.25μ*M*]), 48h for tissue 2 (19.9, 16.5-23.6; [1.25, 0.625]), 144h for tissue 3 (15.2, 3.9-30.4; [0.625, 1.25]), 144h for tissue 4 (6.4, −3.6 to 18.0; [0.625, 0.625]), 48h for tissue 5 (10.1, 8.9-12.5; [0.625, 1.25]), 96h for tissue 6 (24.7, 12.2-36.3; [0.625, 0.625]), 48h for tissue 7 (47.7, 29.6-62.2; [0.625, 1.25]), and 96h for 8 (47.4, 26.8-67.0; [0.625, 0.625]).

**Conclusion:** The concurrent exposure of DEET and permethrin can lead to synergistic cytotoxicity in sinonasal epithelia. Further research is warranted in preclinical animal models to explore whether this synergy accelerates the pathogenesis of chronic rhinosinusitis.

## INTRODUCTION

Pesticides are widely used in agricultural, industrial, and residential settings as well as in the military. Despite their utility in pest control, growing concerns have arisen regarding their potential adverse effects on human health.^1^ Of particular concern is their application as sprays, which facilitates easy inhalation and has led to heightened interest in investigating their relationship with respiratory pathologies.^2–4^ Prior studies have associated pesticide exposure with lower respiratory conditions such as asthma and chronic obstructive pulmonary disease.^1,3^ Associations with upper respiratory conditions such as allergic rhinitis have also been reported.^2,5^

Another upper respiratory condition which has garnered investigational traction is chronic rhinosinusitis (CRS), a persistent inflammatory state of the sinuses characterized by facial pain, nasal obstruction, diminished olfaction, anddrainage.^6^ The most recent 2023 cohort study revealed that residing within 2000 meters of any pesticide application site was associated with a 2.5-fold increase in odds of having CRS.^7^ Delving into the cellular mechanisms underlying this observation, the effects of two common pesticide ingredients, permethrin and *N,N-Diethyl-MetaToluamide* (DEET), on sinonasal epithelial cells (SNECs) were studied in in-vitro exposure experiments. These investigations revealed the cytotoxic effects of both ingredients on SNECs, with exposed specimens exhibiting cell death, morphological changes, and oxidative stress in a dose-response fashion.^8,9^

While understanding the cytotoxic effects of individual pesticide components is important, real-world scenarios often involve simultaneous exposure to multiple ingredients. As such, investigating potential interactions between pesticide ingredients can provide further insights into the practical implications of pesticide-induced toxicity. We present the first investigation into the presence of synergy between DEET and permethrin’s cytotoxicity on SNECs, aiming to further contextualize the potential pathophysiological basis linking pesticide exposure to CRS.

## MATERIALS AND METHODS

### Study Design

SNECs from the ethmoid sinus were procured during endoscopic sinus surgery (ESS) and subsequently exposed to various concentrations of DEET and permethrin, independently and concurrently. Cell viability was monitored using cell growth imaging system during experiments, and subsequent analyses assessed for synergy between DEET and permethrin’s cytotoxicity. Study approval was obtained from the Greater Los Angeles Department of Veterans Affairs Institutional Review Board.

### Cell Culture and Viability Experiments

Sinonasal Epithelial Cells (SNEC) culture system was established in our lab as previously described.^8–10^ The primary monolayer cell cultures of SNECs were used to maintain the homogeneity of the culture system. SNECs were cultured in 96-well plates and exposed to varying concentrations of permethrin (0-5 μM) and DEET (0-5 μM), independently and concurrently, for 156 hours. Dosing concentrations and exposure schedules were based on previously established protocols.^11–13^ Cell proliferation and density were monitored with real-time kinetic data using the *Incucyte* real-time Live Cell Imaging System (*Essen Bioscience*) and phase contrast images.^8,9^ The wells were scanned, and data recorded every 2 hours during the exposure period. Cell density was normalized using the starting level, and data were presented as fold changes since initiation (hour 0) to specific time points during the assay.

### Synergy Calculation

Synergy represents the phenomenon where the combined effects of two agents exceed the expected effects based on a theoretical null model. The null model assumes no interaction between the agents and is often approximated using the Loewe or Bliss additivity equations. The Loewe equation offers an advantage over the Bliss equation as it eliminates the potential for an agent to exhibit synergy with itself during calculation.^14,15^ The Loewe additivity equation operates under the premise of dose equivalence, allowing for the exchange of effects between agents at adjusted-equivalent doses.^16,17^ For example, if the cytotoxic effect of agent A at 2*M* is equivalent to that of agent B at 4*M*, a combination dose of 1*M* of agent A and 2*M* of agent B should yield the same cytotoxic effect.

Synergy is quantified by measuring the degree to which the observed combined effect deviates from the predicted effect according to the null model. When the observed combined effect surpasses the predicted value from the null model, it indicates synergy. If the observed effect is lower than predicted, it signifies antagonism. When the observed effect closely aligns with the predicted value, it suggests additivity. To quantify the synergy between DEET and permethrin, we utilized the *SynergyFinder* software and applied the Loewe additivity model to derive a synergy score (*S_Loewe_*). A score of 0 indicates perfect Loewe additivity; a positive score indicates synergy; and a negative score indicates antagonism.^18^

## RESULTS

The SNECs of 8 patients were procured. Colorimetric assays of cell viability when exposed to various concentrations of DEET and permethrin, individually and concurrently, are illustrated in **Figure 1** (tissue 1-4) and **Figure 2** (tissue 5-8). Loewe synergy scores at 48h, 96h, and 144h when exposed to different-dose combinations of DEET and permethrin are presented in **Figure 3** (tissue 1-4) and **Figure 4** (tissue 5-8).

**Figure 1.**
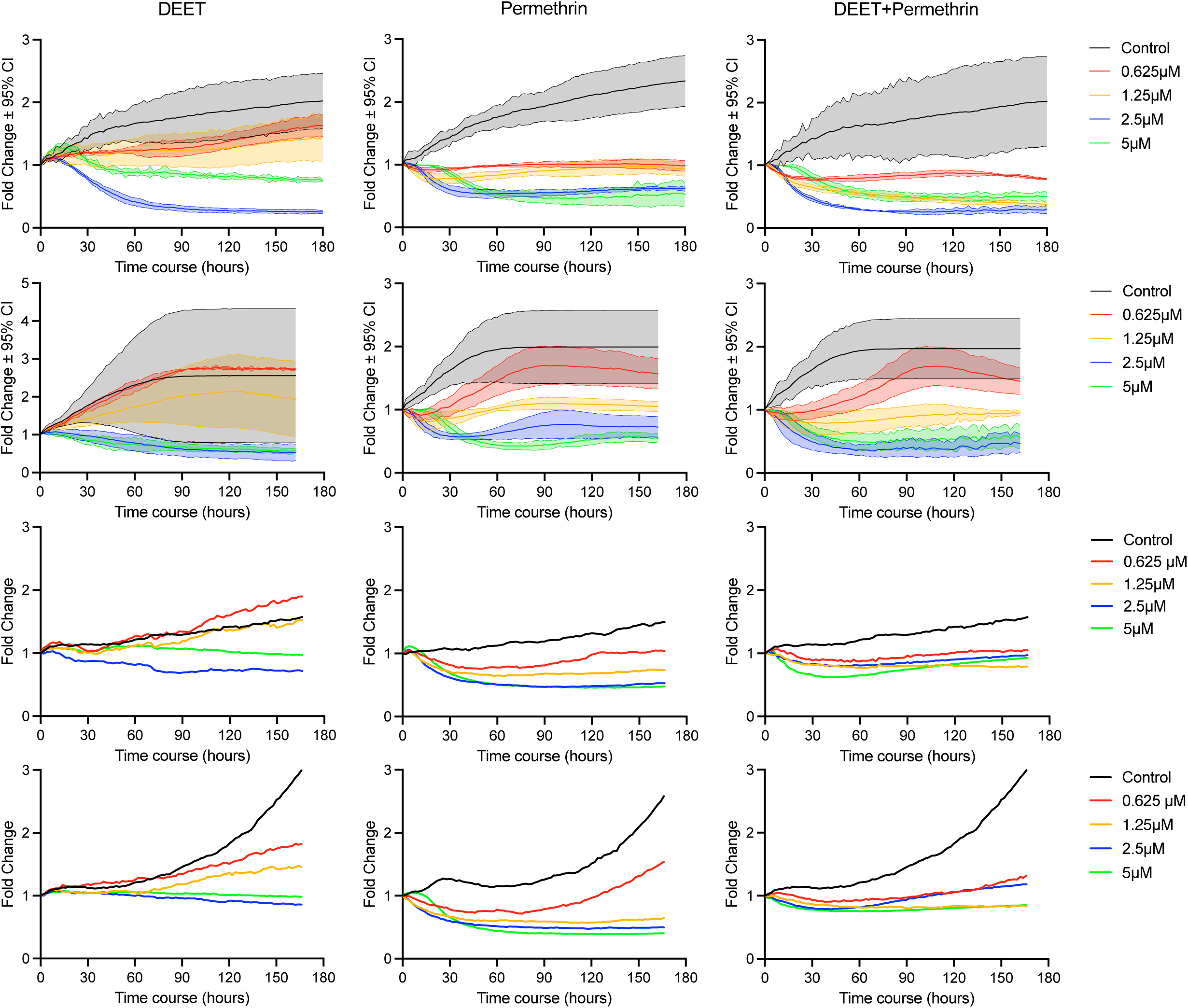
Cell viability of tissues 1, 2, 3, and 4 (top to bottom) when exposed to different concentrations of DEET and Permethrin, independently and concurrently. Confidence intervals are not illustrated for tissues 3 and 4 to enhance clarity of the illustration.

**Figure 2.**
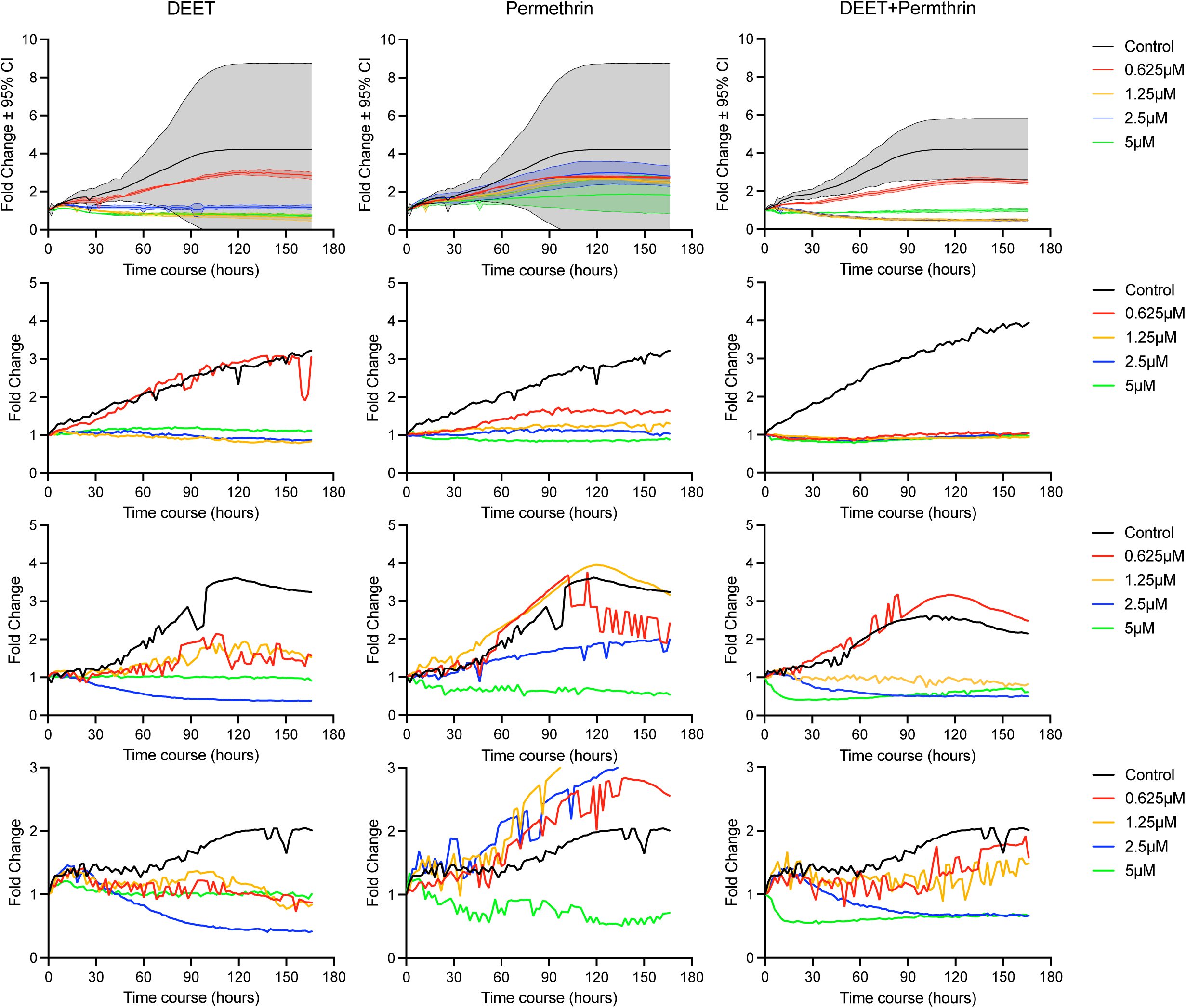
Cell viability of tissues 5, 6, 7, and 8 (top to bottom) when exposed to different concentrations of DEET and Permethrin, independently and concurrently. Confidence intervals are not illustrated for tissues 6-8 to enhance clarity of the illustration.

**Figure 3.**
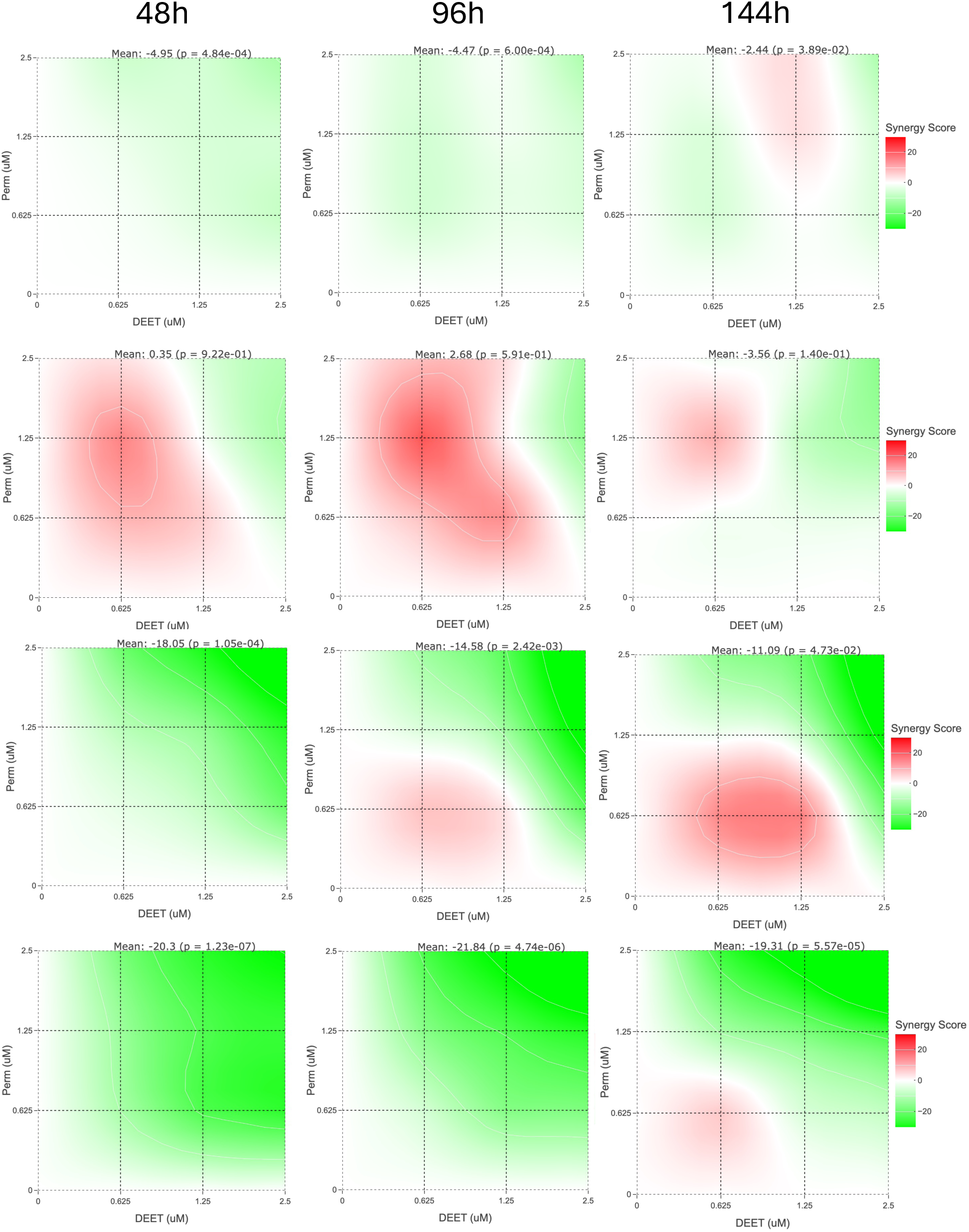
Loewe synergy scores of tissues 1, 2, 3, and 4 (top to bottom) when exposed to different concentrations of DEET and Permethrin.

**Figure 4.**
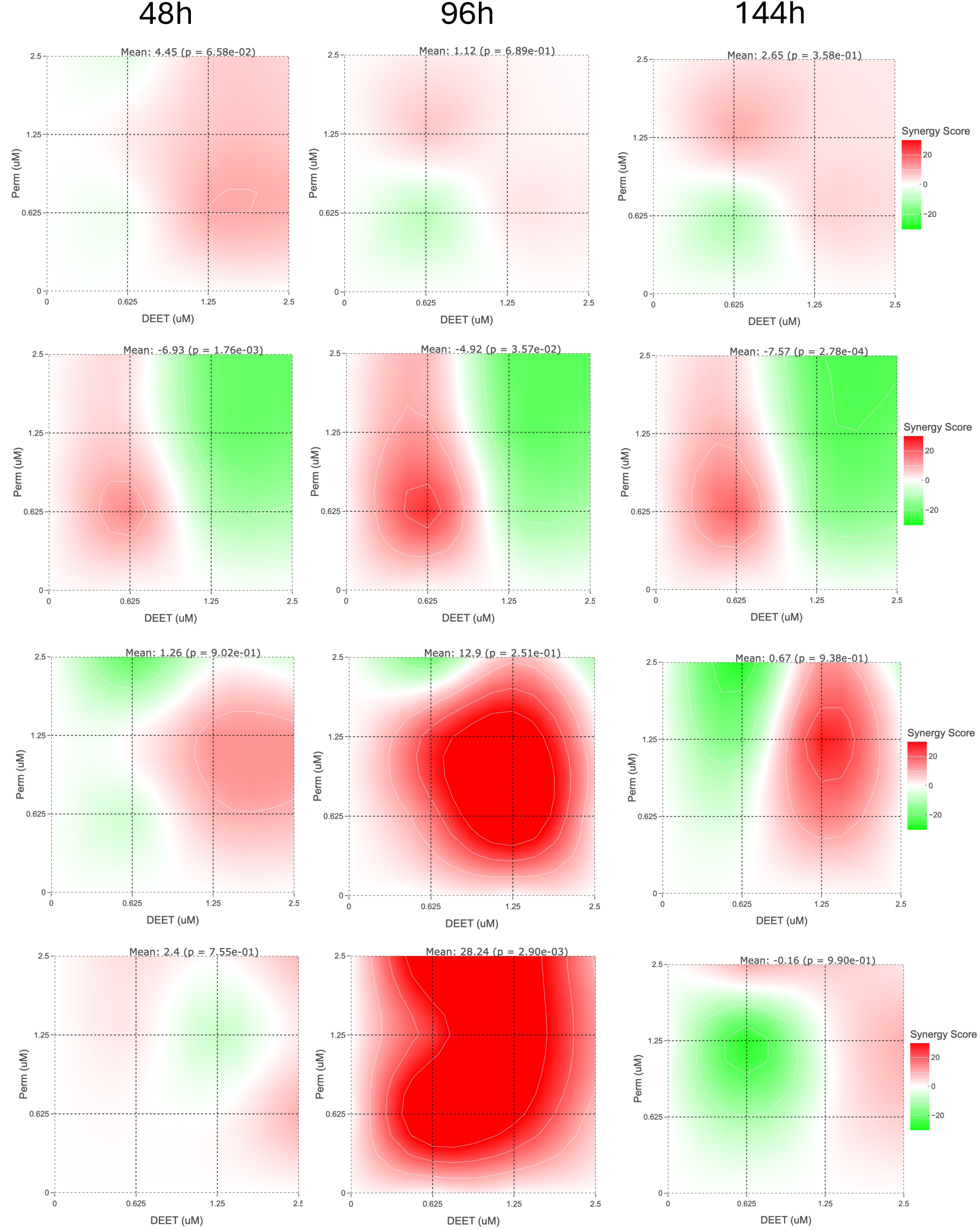
Loewe synergy scores of tissues 5, 6, 7, and 8 (top to bottom) when exposed to different concentrations of DEET and Permethrin.

Tissue 1 was procured from a 76-year old male without a history of CRS. Permethrin demonstrated greater cytotoxicity compared to DEET. At 0.625 μM, cell viability notably decreased with permethrin exposure, whereas only a mild decline was observed with DEET exposure. By 2.5 μ*M*, most SNECs were non-viable in both the DEET and permethrin experiments (**Figure 1** top row). Notably, antagonism was observed until 144 hours, followed by significant synergy at 2.5 μ*M* permethrin + 1.25 μ*M* DEET (*S_Loewe_* 4.15, 95% CI 1.84-7.26) (**Figure 3** top row).

Similar patterns were observed in tissues 2, 3, and 4. Permethrin exhibited greater cytotoxicity than DEET at 0.625 μ*M* in all cases, and most SNECs were non-viable by 2.5 μ*M* in both DEET and permethrin experiments (**Figure 1)**. Significant synergy was observed at various time points and concentrations across tissues, with peak synergy typically occurring between 96 to 144 hours (**Figure 3)**. For tissue 2, significant synergy was already present at 48h, with peak synergy at 96h when exposed to 1.25 μ*M* permethrin + 0.625 μ*M* DEET (*S_Loewe_* 19.87, 95% CI 16.45-23.61) **(Figure 3** second row). Tissue 3 was procured from a 75-year-old-male with CRSwNP. Significant synergy was present starting at 96h, with peak synergy at 144h when exposed to 0.625 μ*M* permethrin + 1.25 μ*M* DEET (*S_Loewe_* 15.18, 95% CI 3.88-30.38) **(Figure 3** third row). Tissue 4 was procured from a 61-year-old-male with CRSwNP. Mild antagonism was present until 144h, where mild synergy was present when exposed to 0.625 μ*M* permethrin + 0.625 μ*M* DEET (*S_Loewe_* 6.42, 95% CI −3.60 to 18.00), (**Figure 3** bottom row).

At 0.625μM, permethrin exhibited greater cytotoxicity than DEET in tissues 5 and 6. However, DEET exhibited greater cytotoxicity than permethrin in tissues 7 and 8, (**Figure 2)**. Significant synergy between DEET and permethrin was evident as early as 48h for tissues 5, 6, 7, and 8 (**Figure 4)**. Tissue 5 was procured from a 41-year-old-male with CRSwNP. Significant synergy was already present at 48h when exposed to higher DEET concentrations, with peak synergy at 48h when exposed to 0.625 μ*M* permethrin + 1.25 μ*M* DEET (*S_Loewe_* 10.11, 95% CI 8.86-12.46) **(Figure 4** top row). Tissue 6 was procured from a 74-year-old-male with CRSsNP. Significant synergy was already present at 48h with lower DEET concentrations, with peak synergy at 96h when exposed to 0.625 μ*M* permethrin + 0.625 μ*M* DEET (*S_Loewe_* 24.65, 95% CI 12.18-36.30) **(Figure 4** second row). Tissue 7 was procured from a 77-year-old-male with CRSsNP. Significant synergy was already present at 48h with higher DEET concentrations, with peak synergy at 96h when exposed to 0.625 μ*M* permethrin + 1.25 μ*M* DEET (*S_Loewe_* 47.68, 95% CI 29.57-62.15) **(Figure 4** third row). Finally, tissue 8 was procured from a 61-year-old-male with CRSwNP. Significant synergy was already present at 48h, with peak synergy at 96h when exposed to 0.625 μ*M* permethrin + 0.625 μ*M* DEET (*S_Loewe_* 47.41, 95% CI 26.76-66.98) **(Figure 4** bottom row).

## DISCUSSION

In this study, we exposed procured SNECs from the ethmoid sinuses of eight patients with varying concentrations of DEET and permethrin. Our findings revealed that permethrin exhibited greater cytotoxicity compared to DEET in most cases. Moreover, concentrations ranging from 1.25 to 2.5 μ*M* of either compound resulted in non-viability in 6 out of 8 of SNECs. Nonetheless, significant synergy in cytotoxicity between DEET and permethrin was observed across all tissues, although the onset and magnitude of synergy varied depending on the specific timepoints and concentrations.

DEET is a widely used insect repellent often applied by spraying or rubbing onto the skin. It has been used in both military and civilian settings.^19^ During the 1991 Gulf War, 75% active ingredient DEET was used by many veterans who later developed a chronic multi-symptom disorder called Gulf War Illness.^20,21^ At the molecular level, DEET has been implicated in carcinogenesis, oxidative stress, germline dysfunction, and immunosuppression.^22–24^ In animal studies, DEET has demonstrated neurotoxicity and pro-neuroinflammatory properties in mouse models, as well as cytotoxicity towards the lysosomal membrane of *Perna perna* mussels.^12,25–28^ Human exposure to DEET has also been linked to elevated risks of encephalopathy.^29,30^ On the other hand, permethrin, a pyrethroid, directly induces neurotoxicity by prolonging the opening of sodium channels, resulting in sustained depolarization, impaired signal propagation, and paralysis.^31^ Although permethrin exhibits selective toxicity towards insects, it is also recognized for its toxicity to human tissues.^32^ Indeed, studies have found permethrin to be nephrotoxic, hepatotoxic, cardiotoxic, immunotoxic, and neurotoxic to humans.^33^ Notably, exposure to permethrin is associated with disrupted hematopoiesis and elevated risks of multiple myeloma.^34–36^ Permethrin was also sprayed liberally on uniforms of military troops during the Gulf War to protect against insect-borne illnesses. Recent research performed with blood samples from Gulf War veterans and in a permethrin exposed mouse model of GWI showed that a permethrin metabolite 3-phenoxybenzoic acid (3-PBA) was associated with peripheral and CNS adaptive immune responses which could contribute to an autoimmune response and neuroinflammation in veterans with GWI.^37^

Although the upper airway is readily exposed to sprayed pesticides, few studies to date have explored the toxicity of DEET and permethrin in sinonasal tissues. A 2023 cohort study revealed that residential proximity to commercial pesticide application sites was associated with higher risks of CRS.^7^ The scientific plausibility of this finding was further substantiated by in vitro exposure assessments of DEET and permethrin on SNECs. In a 2021 study involving three subjects, exposure of SNECs to DEET resulted in a dose-dependent reduction in cell viability and morphological changes suggestive of cellular necrosis.^8^ Similarly, a 2022 study on permethrin demonstrated that increasing concentrations of in vitro exposure led to reduced viability and increased production of reactive oxygen species.^9^

While assessing the cytotoxic effects of individual pesticide ingredients is important, real-world exposure often involves simultaneous exposure to multiple ingredients. Therefore, understanding potential interactions between different ingredients may provide further insights into the practical implications of pesticide-induced toxicity. To our knowledge, the present study is the first to investigate the interactions between two pesticide ingredients with respect to toxicity in SNECs. Specifically, we explored the interactions between two compounds that are commonly used in conjunction, DEET and permethrin. Concurrent application of DEET on the exposed skin and permethrin on the clothing has been recommended for optimal protection against insects since DEET is vapor-active and permethrin does not evaporate.^38^ However, our findings revealed significant synergistic cytotoxicity when SNEC tissues were simultaneously exposed to DEET and permethrin. As such, the potential risks of upper airway toxicity associated with this combination may warrant reconsideration.

Experiments with *Aedes* mosquito species have demonstrated synergistic insecticidal and repellent effects when DEET and permethrin are used in conjunction to develop long-lasting insecticide and repellent-treated nets. While the exact toxicological mechanism responsible for this synergy remains unclear, it is theorized that the synergy is primarily driven by the versatile and distinct modes of action of DEET and pyrethroids.^39–41^ Although the synergy between DEET and permethrin is advantageous in terms of insecticidality, it can lead to heightened toxicity. Prior experimental research exposing rats to a mixture of DEET and permethrin revealed an increase in the incidence of pubertal abnormalities and diseases affecting the prostate, kidneys, testes, and ovaries in subsequent generations. Furthermore, epimutations at DNA methylation regions were observed, indicating potential association with transgenerational disease inheritance.^42,43^ In a subsequent experiment, it was found that the sub-chronic dermal application of DEET and permethrin to adult rats in combination resulted in widespread neuronal cell death observed in the cerebral cortex, hippocampal formation, and cerebellum.^44^ The findings from animal studies prompted investigations into whether these synergistic effects are applicable to humans.

Additional research sought to investigate the link between pesticide exposure and Gulf War Illness with experiments examining the effects of permethrin, DEET, and pyridostigmine bromide (used as a nerve gas agent prophylactic medication) in animal models.^21,45–47^ Results showed neurological, mood and memory changes akin to those seen in Gulf War Illness to be present when these agents were simultaneously administered.^48,49^ Yet, such manifestations were absent when the agents were administered alone, even at concentrations three times higher than those used in simultaneous exposure.^47^ The researchers also noted brain and behavioral changes when DEET and permethrin or other pesticides were concurrently administered under stress conditions, suggesting that stress-induced disturbance to the blood-brain barrier may mediate the observed synergy.^48–52^ Alternative hypotheses posit that heightened toxicity when multiple agents were administered could be attributed to the depletion of detoxification enzymes.^45–47^ Due to the scarcity of literature on pesticide-induced sinonasal toxicity, the precise mechanism underlying the synergy observed in the present study is unclear. We speculate that mechanisms similar to that proposed for Gulf War Illness may be involved. As the nasal cavity serves as the entry point for inhaled pollutants, the activity of cytochrome P-450 (CYP450) is notably higher than that in the liver, primarily attributed to the fourfold greater concentration of NADPH– CYP450 reductase.^53^ As CYP450 is a vital enzyme in the metabolism of permethrin, we hypothesize that co-administration with DEET could expedite the depletion of CYP450, thereby heightening cytotoxicity.^54^ Ultimately, additional research at the molecular level is necessary to fully elucidate the mechanism underlying the synergistic cytotoxicity of DEET and permethrin on SNECs.

Cytotoxicity plays a pivotal role in inflammation, particularly in the pathogenesis of lower airway conditions.^55–57^ As such, exposure to DEET and permethrin, now recognized for their cytotoxic effects on SNECs, may contribute to the development or exacerbation of CRS.^8,9,58,59^ Moreover, our study demonstrates that this cytotoxicity can be synergistic when exposed simultaneously. Upon validation in epidemiological studies, this synergy underscores the substantial impact pesticide exposure can have on SNEC health and subsequent CRS development or exacerbation. It was previously contended that synergistic neurotoxicity resulting from the combination of DEET and permethrin would only manifest at very high concentrations.^38^ However, our experiments utilized concentrations significantly lower than the reported neurotoxicity-inducing concentrations.^8,9^ As such, the SNECs may have heightened sensitivity to this synergy compared to typical cell lines responsible for neurotoxicity. Ultimately, further investigations are necessary using preclinical animal models to ascertain the concentration threshold at which these agents pose a significant risk for inflammation and subsequent CRS development or exacerbation.

### Limitations

There were notable variabilities across tissues with regard to the onset and concentrations for which synergy was significant. However, due to the nature of our study design, we are unable to attribute these differences to any specific patient or environmental factors. The primary aim of our investigation was to ascertain the presence of synergistic cytotoxicity when SNECs are exposed to DEET and permethrin simultaneously. Future studies employing more deliberate patient recruitment and cohort assessment strategies could offer additional insights into the risk factors associated with susceptibility to this synergy. Despite this limitation, our study represents the first exploration of the synergistic cytotoxicity of DEET and permethrin on SNECs. Findings from our investigation provide valuable insights into the practical implications of pesticide exposure and underscore the need for additional research to explore the epidemiological ramifications of pesticide use.

## CONCLUSIONS

DEET and permethrin, frequently combined for pest control, demonstrated in vitro synergistic cytotoxicity to sinonasal epithelial tissues. This heightened cytotoxicity, coupled with inflammation, could potentially contribute to the development or exacerbation of CRS. Therefore, further investigative efforts in laboratory and epidemiologic settings are necessary to fully characterize this relationship and illustrate its necessity for public health deliberations.

## Acknowledgements

The study was funded by the Department of Defense Grant W81XWH-21-2-0048. The authors of this study have no other funding, conflict of interest, or acknowledgement to declare.

## Notes

### Competing Interest Statement

The authors have declared no competing interest.

